# Recurrent DNA Break Clusters Regulated by Polymerase Theta Are Essential for Replication Stress Induced Copy Number Variation

**DOI:** 10.1101/2025.09.01.673480

**Authors:** Lorenzo Corazzi, Alex Ing, Vivien S. Ionasz, Anna Marx, Nathan Trausch, Sarah Benedetto, Giulia Di Muzio, Boyu Ding, Jana Berlanda, Marco Giaisi, Nina Claudino, Thomas Höfer, Pei-Chi Wei

## Abstract

Copy number variations (CNVs) are a form of genetic alteration strongly implicated in numerous neurological and psychiatric disorders, as well as brain cancer. Replication stress is a common cause of CNVs. Despite the prevailing model that CNVs arise from DNA double strand breaks (DSBs), there has been no assay that directly perturbs presumed DSB sources and measures CNV output. Here, we identified a subset of recurrent DNA break clusters (RDCs) as a causal factor for CNV formation. In murine neural progenitor cells under replication stress, mapping the formation of CNVs revealed their location in RDC regions that contain actively transcribed genes. CRISPR/Cas9-mediated transcriptional suppression abrogated both RDC and CNV formation, but does not alter their replication timing. We found that DNA polymerase theta (Pol θ), a protector against CNV formation, plays a critical but context dependent role upstream of RDC formation. Chemically inhibiting the activity of Pol θ reduced end filling and micro-homology-mediated end joining in XRCC4/P53-deficient cells. Conversely, Pol θ inhibition led to elevated DSB density detection at RDC-containing loci in wild-type neural stem and progenitor cells, suggesting its role in preventing transcription-replication conflicts. Our data identify RDCs as contributors to genomic heterogeneity with plausible downstream effects on brain disorders and malignancy.

## Introduction

Copy Number Variations (CNVs) are a pervasive form of genetic structural variation involving deletions, insertions, or duplications of DNA segments^1^. Somatic CNVs can vary widely in size, from kilobases to entire chromosomal arms. CNVs represent a major source of genetic diversity, impacting both normal phenotypic diversities, and the pathogenesis of various genetic disorders, including cancer ^2–5^.

Understanding the mechanisms behind CNV formation is crucial to obtain a full understanding of genetic variation, and to improve diagnostic and therapeutic strategies for CNV-associated disorders. Multiple lines of evidence show that CNVs are dependent on DNA double-strand breaks (DSBs) ^6,7^, which are often repaired via end-joining pathways involving DNA polymerase theta (Pol θ) activity ^8^. Various mechanisms can lead to DSBs. In the context of pathology, replication stress plays a significant role in cancer and certain neuropsychiatric disorders^9,10^. DNA replication stress often leads to stalled replication forks, which can subsequently collapse into DSBs; these DSBs have been proposed as a source of non-recurrent CNVs^7^. Further investigation into the precise interplay between DNA replication stress, and DSB formation, is therefore essential to fully understand the fundamental origins of CNVs, and the disorders they are responsible for.

A major debate is why CNV hotspots cluster at long, actively transcribed, late replicating genes. DNA polymerase inhibition–induced replication stress preferentially generates CNVs at transcriptionally active loci, implicating transcription–replication collisions in CNV formation^11^. Enhancing transcriptional activity at specific genomic loci has even been shown to advance replication timing in vertebrate cell lines, including human cells^12^. These affected loci are typically late-replicating regions, leading to the hypothesis that delayed DNA replication may underlie CNV susceptibility^11,13^. However, it was also shown that genome-wide transient-inhibition of transcription activity did not alter replication timing in CNV-prone genomic regions^14^, suggesting that while late replication timing and CNV formation often coincide, they are not necessarily causally linked. The precise contribution of transcriptional activity, replication timing, and their interplay in the formation of CNVs remains to be elucidated.

In mammalian cells, a substantial number of long genes are hotspots for recurrent DNA break clusters (RDCs)^15–17^. The majority of these long genes encode proteins involved in neural adhesion and synaptic plasticity, suggesting that RDCs may play a role in regulating neurogenesis^18^. RDCs display transcription and DNA replication conflict features, where all RDCs activate transcribed genes. These breaks arise where transcription and replication collide: DSBs accumulate in regions where the two machines move in opposite directions. The DSB orientation tracks the direction of replication forks, implying formation at stressed forks. When forks converge (“inward-moving” forks), the breaks align head-to-head, a 3-D configuration typically resolved by homologous recombination or end-joining^19^. We hypothesize that this configuration promotes deletions and copy number loss in the RDC-containing genes. Consistent with this idea, around one-third of RDC-containing genomic loci were hotspots for CNVs identified in neurons^15,18^, suggesting that a fraction or RDCs result from the same upstream mechanisms that lead to CNV formation.

The primary objective of this study is to elucidate the role of RDCs in CNV formation in murine neural progenitor cells. First, we observed significant and heterogeneous DNA sequence loss in a subset of RDC-containing genes, all of which were actively transcribed and located in late-replicating regions. Second, we experimentally examined the essential role of transcription activity in RDC formation at two RDC-associated genes, *Ctnna2* and *Nrg3*. We found that abolishing transcription suppressed both RDC and CNV formation without altering their replication timing. Third, we characterized an upstream role of polymerase theta (Pol θ), a key factor in CNV formation ^8^, in RDC DSB accessibility. We found that Pol θ has context-dependent roles in NHEJ proficient versus deficient neural progenitor cells in regulating RDC genesis. These findings underscore the critical importance of RDCs in shaping genome heterogeneity and highlight their potential role in the development of neuropsychiatric disorders and cancer.

## Results

### A fraction of RDC-containing genes is copy number variation hotspots in neural progenitor cells

Our initial goal was to determine whether replication stress causes DNA copy-number changes at recurrent DNA-break clusters (RDCs). We treated mouse embryonic stem (ES) cell-derived neural progenitor cells with aphidicolin (Aph), a DNA polymerase inhibitor commonly used for inducing CNVs^11^ and RDCs^18^ with dimethyl sulfoxide (DMSO) used as a control^18^. We carried out whole genome sequencing (WGS) to determine the effect this condition has on DSB and CNV formation (Figure 1A). In the following experiments, unless otherwise indicated, the neural progenitor cells were deficient in XRCC4 and TP53, a genotype combination that enhances DSB recovery, as described in our previous investigations^16,20^. We performed WGS with an average 180-fold coverage of neural progenitor cell genomes to capture heterogeneous CNV patterns. The genomes of neural progenitor cells were predominantly diploid, with the exception of a chromosome 8 gain that was independent of aphidicolin treatment (Supplementary Figure 1). Comparative read-depth analysis revealed a consistent, genome-wide under-representation of RDC loci in Aph-treated cells. When the log₂(Aph/DMSO) average ratios for all 152 RDCs were rank-ordered, five loci fell at least three standard deviations below a size-matched random distribution (Figures 1B-C). In contrast, there were no increases of even two standard deviations, indicating that aphidicolin leads to losses rather than amplifications at these sites, in line with previous findings^11^. Closer inspection of the most strongly affected genes showed striking concordance between replicates (Figure 1C). The top tier of depleted loci is dominated by very large, neuronally expressed genes such as *Grid2*, *Ctnna2*, *Magi2*, *Ilrap1l* and *Sox5*, which are classical hotspots for replication-associated fragility^11,18^.

**Figure 1.**
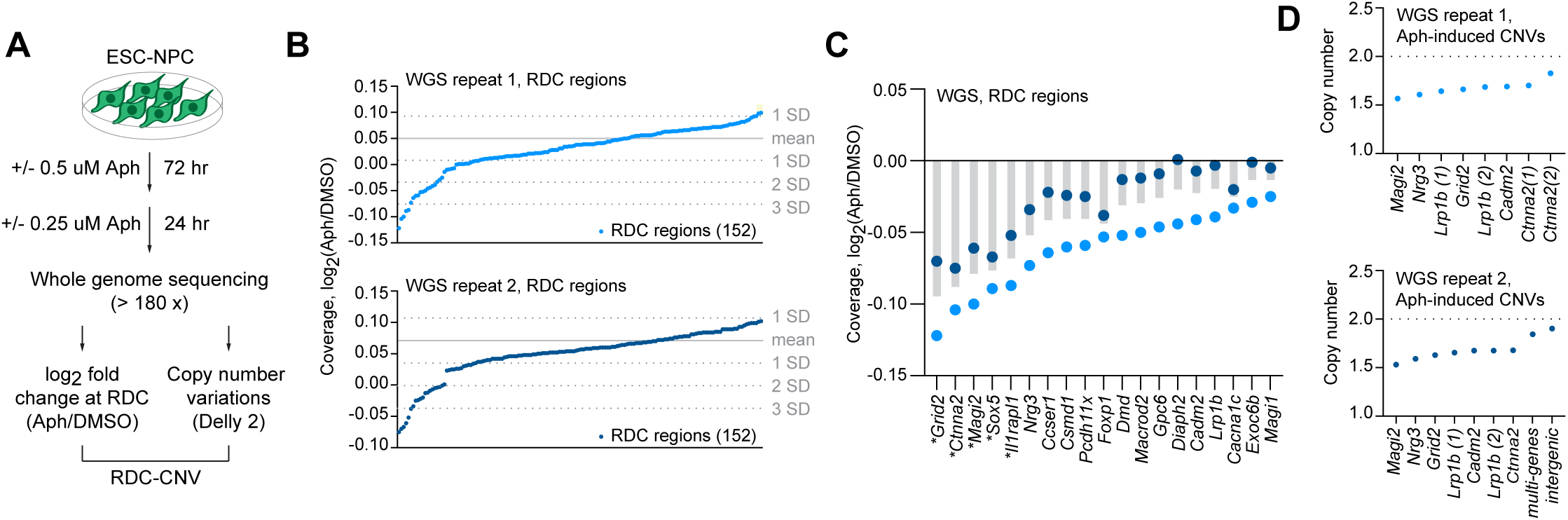
Low-dose aphidicolin elicits shallow copy-number erosion at recurrent DNA-break clusters (RDCs) and discrete heterozygous deletions in ES cell-derived neural progenitors (NPCs). (A) Experimental design. Mouse ESC-derived NPCs were treated with 0.5 µM aphidicolin (Aph) for 72 h followed by 0.25 µM Aph for 24 hours, or with solution (DMSO) for the same period. Genomic DNA from two independent biological replicates was subjected to > 180× whole-genome sequencing (WGS). Read depth was analysed (i) for log₂(APH/DMSO) fold-change across 152 previously defined RDC intervals and (ii) for structural copy-number variants (CNVs) using Delly2; together these outputs are referred to as the RDC-CNV dataset. (B) Rank-ordered log₂(Aph/DMSO) coverage for every RDC in WGS replicate 1 (upper) and replicate 2 (lower). Blue dots, individual RDCs; grey solid line, mean of 1,520 size-matched random regions excluding RDC regions; grey dotted lines, ±1-3 s.d. of the random set. (C) Per-gene view of the 19 RDCs showing the largest coverage losses (left to right; gene symbols on x-axis) in both replicates. Genes that deviate by more than 3 SD in Figure 1B are indicated with a star. Grey bars show the average of replicate 1 (light blue dots) and replicate 2 (dark blue dots) values. (D) Copy number for CNVs detected in replicate 1 (top) and replicate 2 (bottom) by Delly 2. Each dot represents a discrete CNV; gene names denote overlap with an RDC from panel C. The dashed line marks the diploid baseline (2 copies); all events correspond to single-copy losses (median ≈ 1.6 copies).

Structural-variant calling corroborated these findings. Across both datasets, we identified nine discrete CNVs using a variant calling tool Delly2^21^ (Figure 1D and Supplementary Data 1). All exhibiting relative copy numbers of 1.5∼1.8, consistent with heterozygous deletions. Six of these deletions mapped directly within an RDC showing significant DNA sequence losses (*Magi2*, *Nrg3*, *Lrp1b*, *Grid2*, *Cadm2*, and *Ctnna2*). Two CNVs mapped in repeat 2 (one multigenic and one intergenic) were not reproducible and were therefore excluded from the following analysis. Together, these data show that low-dose aphidicolin produces two hierarchies of instability at RDC genes: (i) a pervasive, shallow depletion of sequencing coverage across roughly one-fifth of all RDCs, suggestive of incomplete genome duplication or sub-clonal loss, and (ii) rarer, focal deletions confined to a subset of the most vulnerable loci. The concordance between biological replicates underscores the reproducibility of these events.

### Heterogeneous DNA sequence loss within RDC-CNV hotspots

Neural progenitor cells exposed to aphidicolin revealed discrete copy-number losses in six RDC-containing loci, situated within late <u>C</u>onstant-<u>T</u>iming <u>R</u>egions (CTRs) (Figure 2A). We noted that Delly2 has a cutoff that could not recover all genomic areas showing copy number loss. In some cases, the maximum loss in read depth reaches almost the entire allele (*Ilrapl1*, *Ccser1*; Supplementary Figure 2A). At each locus, the log₂(Aph/DMSO) read-depth ratio dipped by ≈ 0.3– 0.5, consistent with heterozygous deletions spanning 0.3–2 Mb. To precisely map DSBs in genes with CNVs, we re-plotted the RDC DSBs determined by capture-ligation based assay, linear amplification-mediated, high-throughput genome-wide translocation sequencing, LAM-HTGTS^16,22^. This assay mapped unidirectional clusters of single-ended DSBs precisely within these deleted intervals^16^, indicating that aphidicolin-induced fork stalling converts recurrent breakage into segmental loss. Despite the structural change, high-resolution Repli-seq profiles showed only a minor shift in replication timing: the affected regions remained uniformly late-replicating under both solvent and stress conditions. In contrast, we did not detect CNVs at RDC regions located within replication timing transition regions (TTRs; Supplementary Figure 2B) or biphasically replicating domains (Supplementary Figure 2C), the latter of which have been associated with common fragile site formation^23^.

**Figure 2.**
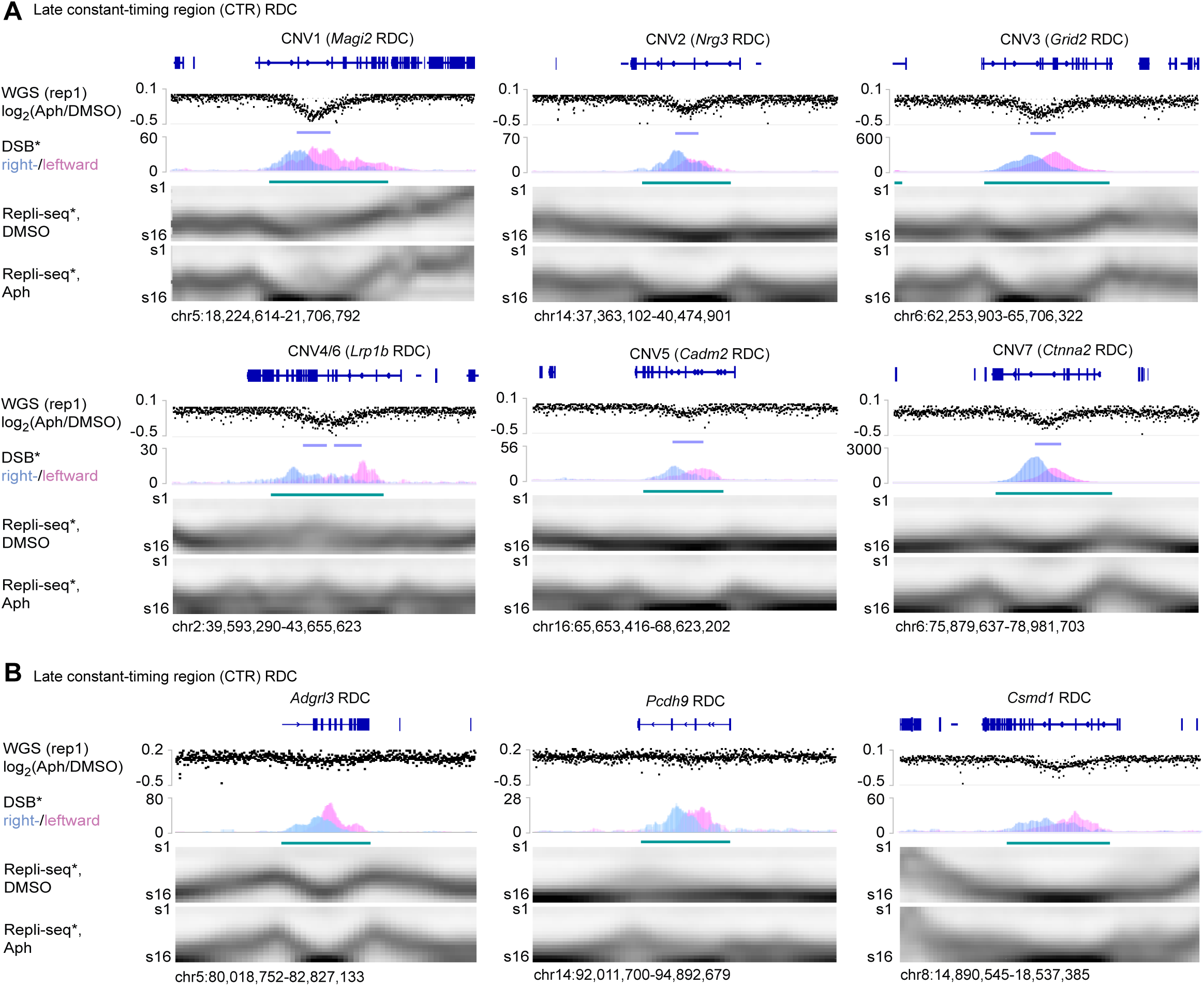
Aphidicolin-induced heterozygous deletions arise at a subset of late-replicating constant-timing recurrent DNA break clusters. (**A)** Six copy-number variants (CNV1–CNV7) identified in neural progenitor cells after 96 h of aphidicolin (Aph) treatment. For each locus (gene names indicated above), the upper track shows the whole-genome-sequencing log₂(Aph/DMSO) coverage ratio (WGS, replicate 1); downward deflections mark heterozygous losses (turquoise bars). The middle track plots strand-specific LAM-HTGTS signal for single-ended double-strand breaks (DSB), coloured by right-(blue) and left-ward (pink) polarity; shaded distributions locate clustered breaks that coincide with CNV boundaries. The lower two heat maps present Repli-Seq replication-timing profiles for solvent control (DMSO) and APH-treated cells (S-phase fractions S1– S16), illustrating that each CNV lies within a late constant-timing region (CTR) whose timing is unchanged by Aph. Data were extracted from a previous study^16^ and re-plotted. Genomic coordinates are given beneath each panel. (**B)** Three additional late CTR RDC hotspots (*Adgrl3, Pcdh9, Csmd1*) accumulate Aph-induced DSB clusters below Delly2 significance cutoff. *Adgrl3* and *Pcdh9* did not show significant DNA copy number loss, while *Csmd1* showed significant DNA sequence losses (Figure 1C). Tracks are as in (A).

Not all RDC loci at the late CTR followed this trajectory. Three other late CTR loci (*Adgrl3*, *Pcdh9*, and *Csmd1*) accumulated equally robust break clusters after aphidicolin yet did not suffer significant copy number loss (Figure 2B). To test if this difference was contributed by under replication, we calculated underreplication indexes (URI)^14^ for RDCs located in the late CTR. Comparing Aph-treated to solution-treated (DMSO) cells, we found that not all CNVs correlated with under-replication, defined by low URI scores (Supplementary Figure 2D). In addition, we also observed cases where low URI at the *Csmd1* and *Nkain2* loci did not account for CNV formation, and vice versa (Supplementary Figure 2D), suggesting that these differences were not solely due to incomplete DNA replication.

These observations indicate that replication-stress-induced breaks at fragile sites are not entirely replication timing-dependent, and can be resolved through locus-specific pathways.

### RDC-dependent CNV formation is not associated with alterations in replication timing

To test whether transcription of large neuronal genes is required for the formation of RDCs, we used CRISPR–Cas9 to delete the proximal promoter/enhancer of *Ctnna2* and *Nrg3* genes (Figure 3A). Two independent neural progenitor cell clones derived from ES cell lines (pe-del1 – 4; Figure 3B) were used for each gene locus, and the unique deletion was validated by Sanger sequencing (Supplementary Figure 3). By mapping nascent RNA with the global run-on sequencing (GRO-seq) assay^24^, we found that the nascent transcripts naturally encoded at the minus strand in the parental cells disappeared in the promoter/enhancer-deleted ES cell-derived neural progenitor cells (Figure 3B, green panels). Mapping DSBs in the same cell in parallel experiments showed that RDC clusters vanished from the *Ctnna2* and *Nrg3* locus in promoter/enhancer-deleted ES cell-derived neural progenitor cells (Figure 3B, DSB panels). This data strongly indicates that RDCs at *Ctnna2* and *Nrg3* were transcription-dependent.

**Figure 3.**
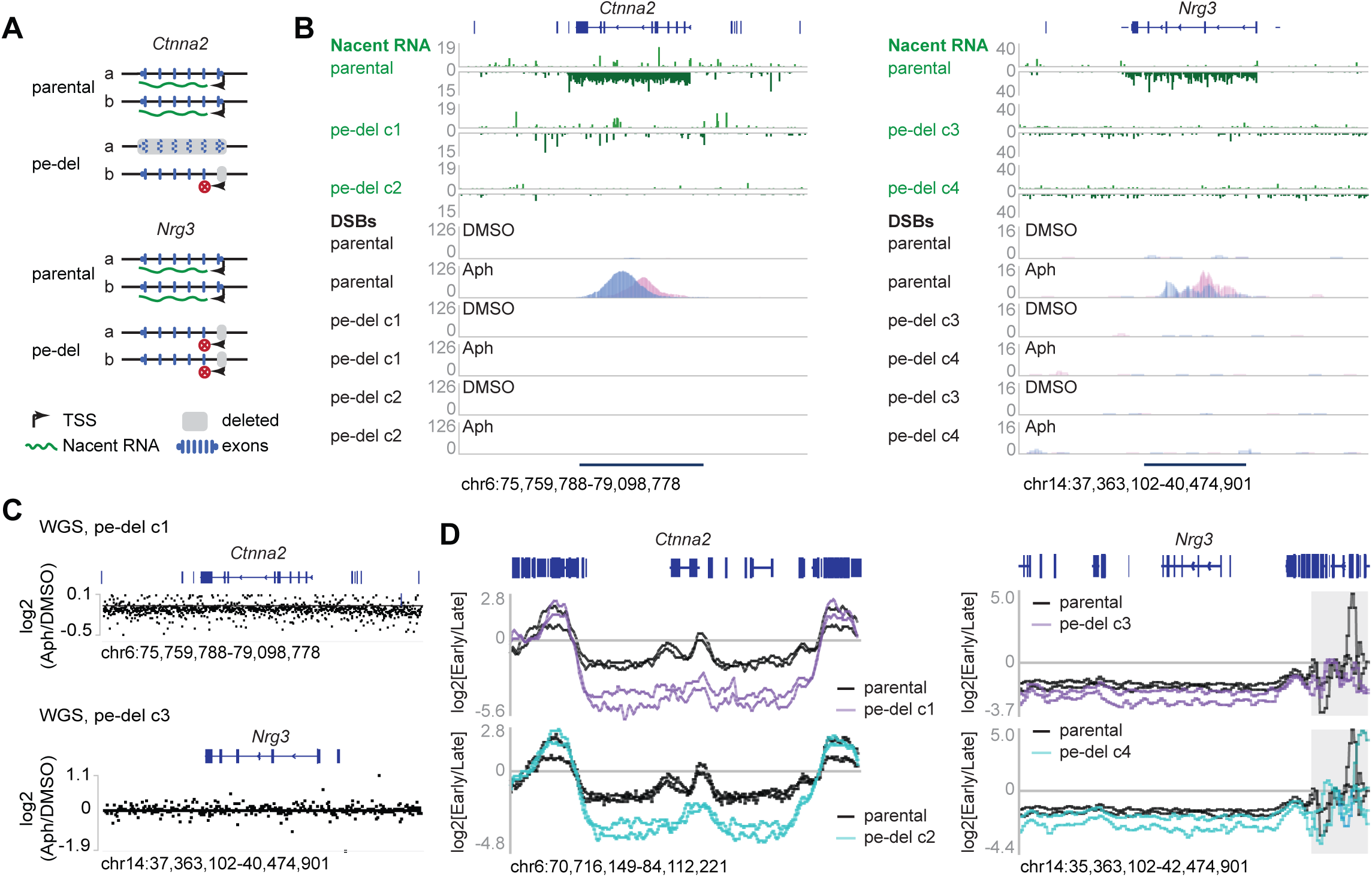
Promoter/enhancer deletions extinguish transcription-dependent RDCs at the CNV-containing genes *Ctnna2* and *Nrg3* without elevating their replication timing. (**A**) Strategy for CRISPR/Cas9 deletion of the major promoter/enhancer (pe-del) that drives the long neuronal isoform “a” of the giant genes *Ctnna2* (left) and *Nrg3* (right). Grey boxes mark the region removed in two independent clones (c1, c2); red X markers indicate transcription inactivation. Isoform-specific transcription start sites (TSS) and exons are depicted below. (**B)** Loss of nascent transcription eliminates RDCs. Data (green) shows gene-body-spanning transcription of *Ctnna2* (left) and *Nrg3* (right) in parental cells, which is almost completely abolished in both pe-del clones. LAM-HTGTS tracks (blue/pink, right/left-ward single-ended DSBs) reveal a sharp, aphidicolin-induced DSB peak in parental cells that disappears in clones with deleted promoters. Scale bars: DSB per thousand. (**C)** WGS log₂(Aph/DMSO) coverage ratio around the *Ctnna2* locus in the *Ctnna2* promoter/enhancer-deleted cells (pe-del c1, upper), and around the *Nrg3* locus in the *Nrg3* promoter/enhancer-deleted neural progenitor cells (pe-del c3). (**D)** Replication-timing (Repli-seq) profiles across the two loci. Log₂(Early/Late) ratios are plotted for parental cells (black) and the two pe-del clones in untreated conditions. The grey box at the *Nrg3* adjacent region denotes a repetitive sequence-rich area.

To test whether DNA sequence losses at CNVs were RDC dependent, we performed high-coverage (average 180×) whole-genome sequencing on neural progenitor cells with promoter/enhancer deletions, both before and after mild aphidicolin treatment (Figure 3C). We observed consistent CNV losses at *Magi2, Ccser1, Grid2, Sox5, Gpc6*, *Prkg1*, and *Lrp1b* CNV in the deleted clones (Supplementary Data 1). Significantly, the log₂(Aph/DMSO) ratio indicated that copy number losses at the *Ctnna2* (pe-del c1) or the *Nrg3* (pe-del c3) loci were no longer present in these clones (Supplementary Data 1). These data show that Aph-induced CNVs at the *Ctnna2* and *Nrg3* genes are dependent on their transcription, further suggesting that these CNVs result from RDCs.

As shown in previous studies^12^, transcription activity can delay replication, resulting in under-replicated genomes that coincide with CNV hotspots. To evaluate DNA replication timing switching, we performed two-fraction replication sequencing (Repli-Seq) on the parental neural progenitor cells, as well as on the promoter/enhanced-deleted cell lines (Figure 3D). At the *Nrg3* locus, transcription ablation did not further alter the replication timing. At the *Ctnna2* locus, the replication of the locus and the adjacent area was even delayed. These data indicated that the loss of CNV from these cells was not the result of advanced replication timing.

Together, our data suggested that RDCs at the *Ctnna2* and *Nrg3* loci are transcription-dependent, and DNA sequence loss in CNV hotspots is RDC-dependent.

### Pol θ promotes RDC accessibility in XRCC4/P53-deficient neural progenitor cells

Next, we sought to determine how polymerase theta (Pol θ) activity affects RDC-derived end availability for joining. It has been suggested that, in the non-homologous end-joining (NHEJ) deficient cells, DNA repair and chromosomal translocations are mediated by Pol θ^25,26^. Specifically, Pol θ fills short gaps at single-ended replication breaks^27^, revealing 1–2-bp microhomologies that drive repair through Pol θ-mediated end-joining (TMEJ). Therefore, inhibition of Pol θ should divert repair toward blunt ligation. Conversely, Pol θ regulates DNA replication in a domain-dependent context^28^, suggesting its role in DNA replication. Moreover, Pol θ promotes structural variant formation at common fragile sites in M phase^8^, indicating a role in CNV formation. Our data suggests that RDCs are required for CNVs at *Ctnna2* and *Nrg3* loci (Figure 3). Although it is plausible that Pol θ assists TMEJ to form CNVs, to our knowledge, it is not clear whether Pol θ regulates DSB end accessibility at RDCs (Figure 4A).

**Figure 4:**
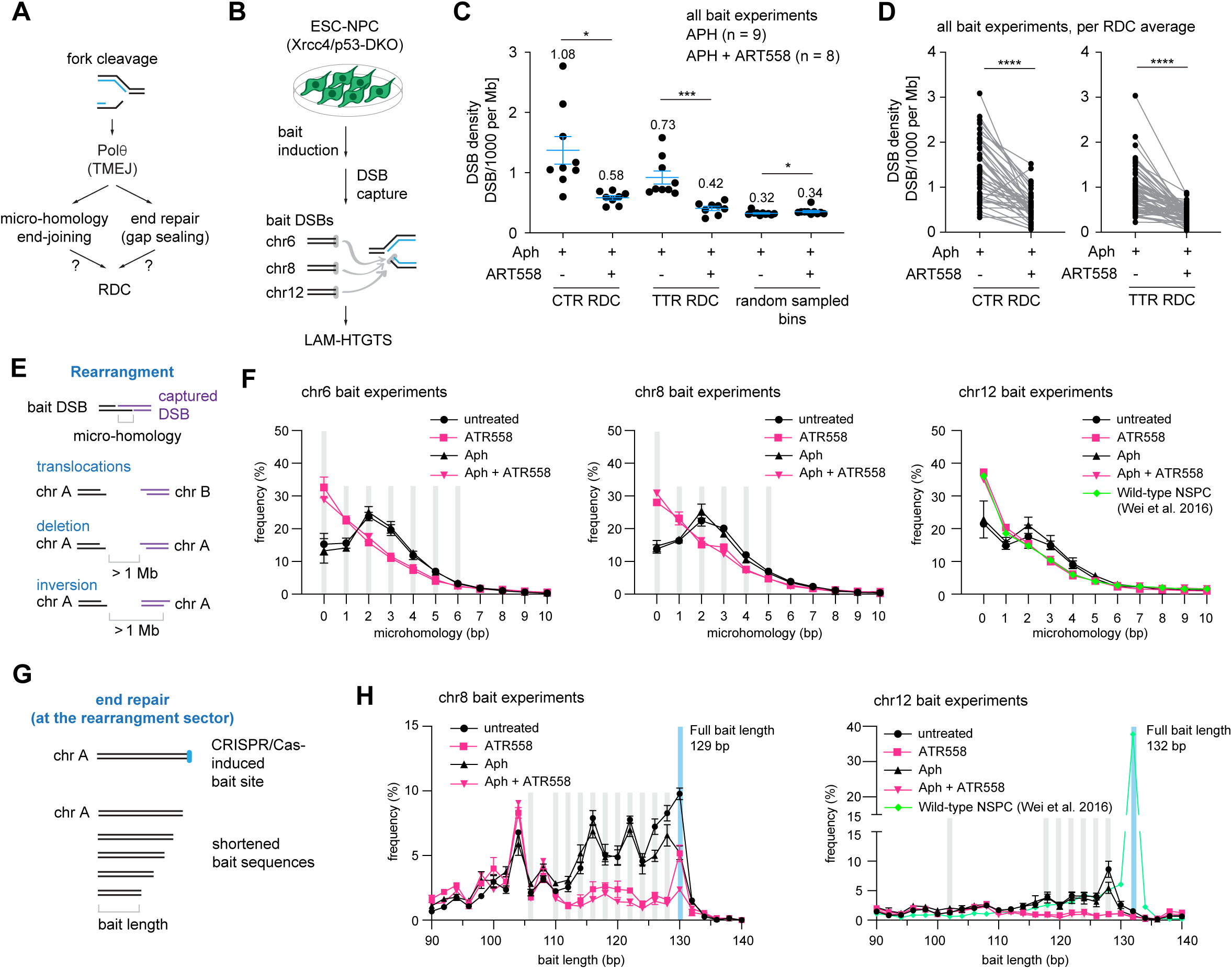
Pol θ activity shapes break formation, capture efficiency and repair pathway choice at RDCs. **(A)** A proposed model for the Pol θ function in RDC formation. At a stalled replication fork, a single-ended single-strand break reprograms the fork structure. Pol θ is proposed to repair the broken end and assist microhomology search, promoting RDC stabilization and theta-mediated end joining (TMEJ) in XRCC4/p53-deficient neural progenitor cells. (**B)** Schematic of the DSB capture experimental design. CRISPR-Cas9 “bait” DSBs were induced on chromosomes 6, 8 and 12 in the XRCC4/p53-deficient neural progenitor cells; cells were then treated with aphidicolin (APH) and/or the Pol θ inhibitor ART558. DSBs were then quantified using LAM-HTGTS. (**C)** DNA break density shown as Junction Per Thousand per Megabase (JPTM) within RDCs at the late CTR (42 RDCs), at the timing-transition region (TTR, 65 RDCs), and randomly shuffled RDC-equivalent size genomic bins (1520 bins). Plot shows the average and SEM of DSBs within the indicated category in Aph-treated (9 experiments) or Aph and ART558-treated (8 experiments) neural progenitor cells. The mean and the standard error for the mean (SEM) were plotted; the mean value for each condition is annotated. Statistical significance was determined using a two-tailed t-test followed by a false recovery rate (FDR) correction. (**D)** Before– and-after plot showing the paired DSB density change per RDC at the CTR region (left) or at the TTR region (right). Dots represent the average DSB density from nine (Aph) or eight (Aph + ART558) independent experiments. Lines connected the same RDC in two conditions. Statistical power was determined using a two-tailed Mann-Whitney paired T-test. (**E)** Schematic defining of rearrangement junction captured when a bait DSB joins a break on the other chromosome (translocation) or on the same chromosome (inversion or deletion) with an endogenous DSB 1 Mb away from the bait. (**F)** Micro-homology frequency at translocation junctions at chromosome 6, 8 and 12 baits. The mean and the SEM are shown. Bins showing statistically significant differences between Aph and Aph + ART558 samples, as determined using a two-tailed t-test, are indicated by grey shading. (**G)** A schematic illustrating the measurement of bait-end resection at translocation junctions. (**H)** Distribution of recovered bait lengths in translocation-sector reads for chromosome 8 and chromosome 12. Mean (dots) and SEM (bars) are shown. Bins exhibiting a statistically significant difference are shaded using the same approach as in panel F. For **(F)** and **(H),** values for individual repeat are shown in Supplementary Tables 2-3; junction numbers analysed per experiment, and statistical significance per bin are shown in Supplementary Table 4.

To assess how Pol θ influences RDC DSB end accessibility, we employed LAM-HTGTS to map DSB junctions across the genome. Briefly, a CRISPR-Cas9 induced “bait” DSB was introduced on chromosomes 6, 8, or 12 in XRCC4/p53-deficient neural progenitor cells. These cells were then left untreated or exposed to ART558, an inhibitor of Pol θ DNA synthesis activity^29^, aphidicolin, or both, before being collected for LAM-HTGTS (Figure 4B). It should be noted that LAM-HTGTS measures the frequency of recoverable (i.e., processed and joinable) ends rather than absolute breakage. As a result, perturbations that alter end processing, for instance, resection/gap-filling and microhomology usage, will change the number of DSBs captured but not necessarily the DNA lesion created.

We found that loss of the polymerase activity of Pol θ reduced the number of preys captured junctions mapping to RDCs (Figures 4C-D), while the number of preys captured remains higher than the Pol θ-inhibition alone at non-RDC genomic bins (Figure 4C). We observed that junction losses were associated with a redistribution of LAM-HTGTS junction clusters towards the center of the RDC loci. For example, at the *Ctnna2* and *Nrg3* loci, the distance between the head-to-head DSB clusters was reduced in aphidicolin-treated cells combined with Pol θ inhibition (Supplementary Figure 4A). Although previous literature has suggested that the effect of Pol θ would be more pronounced at the G2/M phases, we also observed the reduction in RDC-mapped junction recovery in RDCs within TTR regions (Figure 4C). At the *Large1* locus, the rightward-oriented junction cluster was not recovered (Supplementary Figure 4A).

We speculated that inhibiting Pol θ polymerase activity would result in reduced micro-homology dependency and gap filling^27,30^, thereby decreasing DSB end accessibility at RDCs and reducing the number of preys captured by LAM-HTGTS. To test this hypothesis, we analyzed the micro-homology usage at the translocation junction formed between the LAM-HTGTS “bait” and the endogenous “prey” DSB ends. We grouped the junctions into two categories: the junction formed between the “bait” DSB and endogenous DSB lies on a different chromosome (translocations), and a junction formed between the “bait”, and the DSB downstream on the same chromosome (the deletions and inversions) (Figure 4E; Supplementary Data 2,4). In Xrcc4/p53-deficient cells, as expected, we observed a preference of two base pair micro-homology usage at the rearrangement junctions (Figure 4F). In contrast, Pol θ inhibition reverted the micro-homology usage to direct joining, even equivalent to the level observed in the wild-type cells (Figure 4F). This is also true for RDC-containing genomic loci (Supplementary Figure 4C). Hence, the gap filling defect affects DSB ends genome-wide, including at RDCs.

To examine whether the loss of micro-homology is linked to end processing – including resection and gap filling, we analyzed the total length of the translocated DNA sequence (Figures 4G, H; Supplementary Figure 4D; Supplementary Data 3,4). This analysis was limited to the “bait” compartment, as the origin of “prey” prior to end processing was known. For both rearrangements and deletions around the bait sites, we observed a significant and reproducible reduction in the representation of bait length, with an average shortening of approximately 10-15 base pairs. This observation suggested that the polymerase activity of Pol θ either protects the DNA ends from extensive resection or is required for filling the single-strand DNA gap, thereby creating end-joining-ready DNA breaks.

In summary, RDC derived end accessibility to translocation capture by LAM-HTGTS is Pol θ polymerase activity-dependent, mediated by micro-homology-dependent joining, and single-strand DNA filling in XRCC4/p53-deficient neural progenitor cells.

### Pol θ inhibits RDC-derived junction capture in wild-type neural stem and progenitor cells

The role of Pol θ as an antagonist of RDC derived junction capture in the XRCC4/p53-deficient context may be due to their addition to theta-mediated end joining, in NHEJ-deficient conditions^31^. To investigate the role of Pol θ in the XRCC4-proficient condition, we isolated primary neural stem and progenitor cells from the wild-type mouse brains at embryonic day 17.5. We transfected the CRISPR-Cas9-expressing vector into the neural stem and progenitor cells at 10 days *in vitro*, followed by aphidicolin treatment and Pol θ inhibition, in the same manner as the XRCC4/p53-deficient neural progenitor cells (Figure 5A).

**Figure 5.**
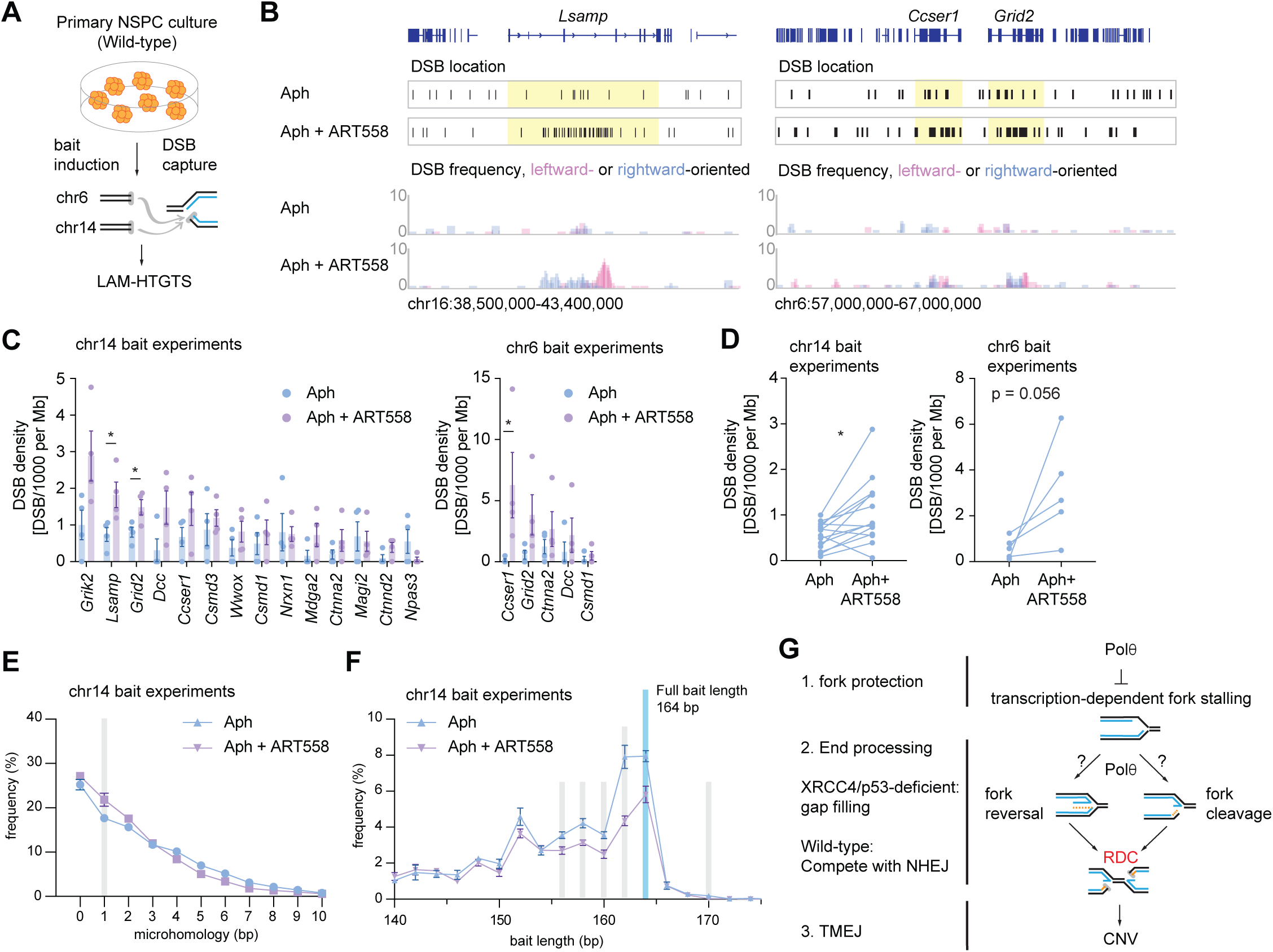
Pol θ inhibition enhances DSB density at the RDC loci. **(A)** Experimental diagram. **(B)** DSB distribution at the *Lsamp* and *Ccser1-Grid2* loci. The exact DSB location and DSB frequencies are shown in separate panels. The corresponding gene loci on the DSB location tracks are highlighted in yellow. 5,908 and 6,071 total interchromosomal junctions were plotted for APH and Aph plus ATR558 treated conditions, respectively. **(C)** DSB density shown as Junction Per Thousand per Megabase (JPKM) for the RDC hotspots identified in wild-type NSPCs. The figure shows the mean and SEM. Statistical significance was obtained using a two-tailed t-test. The figure shows the mean and SEM. *: P <0.05. Hotspots identified in the chr14 and chr6 experiments respectively, are shown here. **(D)** Paired DSB density change per RDC in the chr14 and chr6 experiments, respectively. Statistical significance was determined by a two-tailed Mann-Whitney paired T-test. * P <0.05. **(E)** Micro**-**homology frequency at translocation junctions at chromosome 14 baits. The figure shows the mean and SEM. **(F)** Distribution of recovered bait lengths in deletion-sector reads for chromosome 14. The figure shows the mean (dots) and SEM (error bars). For **(E)** and **(F),** values for individual repeat are shown in Supplementary Tables 2-3; junction numbers analysed per experiment, and statistical significance per bin are shown in Supplementary Table 4. **(G)** A working model for RDC-dependent CNVs. In neural progenitor cells, CNV formation depends on transcription-mediated RDCs.

We performed LAM-HTGTS to analyze the DSBs captured by two chromosomal baits, and to assess the DSB junction recovery mapped to RDCs previously identified in wild-type neural stem and progenitor cells^18^. At the *Lsamp* gene locus, the most promising RDC in wild-type neural progenitor cells^18^, we observed a significant increase in RDC-derived junction recovery upon Pol θ inhibition (Figure 5B). Junction gain was also true for the *Grid2* and *Ccser1* loci (Figure 5B), suggesting a global increase in RDC-mapped junction recovery at RDC hotspots. Indeed, all but two RDCs with multiple recovered junctions exhibited significantly enhanced DSB junction detection upon Pol θ inhibition (Figure 5C, D), suggesting that polymerase theta competes with NHEJ to repair DSB ends formed in RDC regions.

We further investigated the micro-homology usage and resection status in wild-type neural stem and progenitor cells. In contrast to the XRCC4/p53-deficient neural progenitor cells, wild-type neural stem and progenitor cells were less dependent on theta-mediated joining (Figure 5E and Supplementary Figure 5A; Supplementary Data 2,4). Analyzing the total length of LAM-HTGTS bait DNA sequences revealed a slight reduction in the bait length (Figure 5F and Supplementary Figure 5B; Supplementary Data 3,4), much less pronounced than that in XRCC4/p53-deficient neural progenitor cells. In summary, these results suggest that Pol θ activity suppresses the end-repair mechanisms that lead to an extension in DNA break ends. The role of Pol θ in regulating DSB accessibility is context-dependent with respect to RDC formation. In wild-type cells, Pol θ suppresses RDC-prone hotspots, potentially by modulating replication timing^28^, or through competition with NHEJ. In the absence of non-homologous end joining, neural progenitor cells depend on Pol θ for end processing and the facilitation of microhomology-mediated end joining.

## Discussion

### Summary of observations

In this study, we establish that transcription-dependent RDCs generated under replication stress directly cause somatic CNVs at ultra-long, late-replicating neuronal genes. Using deep whole-genome sequencing, CRISPR/Cas-mediated transcriptional silencing, and targeted manipulation of Pol θ, we trace the causal trajectory from transcription-replication conflict, to DSB and subsequently CNV formation. Replication stress induced by low-dose aphidicolin converts a subset of ultra-long, late-replicating neuronal genes that harbour RDCs into CNV loss hotspots. Silencing *Ctnna2* and *Nrg3* promoters abolishes both transcription-dependent RDCs, and the associated CNVs, without altering replication timing, indicating that sustained transcription, rather than replication lateness, drives breakage and loss. Pol θ then shapes lesion fate: in XRCC4-deficient cells, its inhibition suppresses capture of aphidicolin-induced breaks by blocking microhomology-mediated gap filling, whereas in wild-type progenitors, the same treatment increases breakage at RDC hotspots, implying a fork-protective role. Together, these observations support a model in which transcription-driven RDCs initiate somatic CNVs, with Pol θ acting as a context-dependent modulator of break formation, processing, and repair (Figure 5G).

### Transcription-dependent RDCs are a prerequisite for the formation of CNVs

We found that CNVs at *Ctnna2* and *Nrg3* are transcription-dependent (Figure 3), consistent with our hypothesis that S-phase transcription may lead to transcription-replication collisions, causing persistent fork stalling and single-stranded DNA intermediates^16^. It is known that transcription-coupled nucleotide excision repair can leave short ssDNA gaps as repair intermediates^32^. In parallel, topoisomerase activities required for long-gene transcription can introduce ssDNA lesions: topoisomerase I (Top1) processing at embedded ribonucleotides generates a nick and, when trapped as a cleavage complex, Top1cc, a second vulnerable site; When replication-forks encounter these gaps they are converted to single-ended DSBs^33^. The gap-to-break mechanism has primarily been described in early-replicating regions of the genome^34,35;^ the transcriptional dependent-mechanism of RDCs in late-replicating domains warrants further investigation. We speculate that RDC locus accessibility may interfere with the conversion of RDCs to CNVs. Long and late-replicating genes, and heterochromatin, are in spatial proximity with the lamina-associated domains (LADs)^36^. It has been suggested that LADs belong to the B compartment^37^, where the close spatial proximity of DNA sequences within a given LAD may enhance the efficiency of end searching required for end joining. In this regard, analyzing RDCs at late CTR revealed that five out of six genes located within CNVs appeared to associate with LADs (Supplementary Figure 6). On the other hand, two of the CNV-free, strong RDC-containing loci (*Adgrl3* and *Pcdh9*) are not associated with LADs. In total, for late-replicating RDCs that do not express CNVs, only 17 of 57 RDCs are associated with LADs. In this regard, CNV-containing RDCs were significantly enriched with LADs (chi square test; p = 0.009). Yet, how LADs regulate CNV formation remains to be clarified.

### The polymerase activity of Pol θ promotes RDC end accessibility in the context of XRCC4/p53-deficiency

NHEJ-deficient cells depend on Pol θ to repair DSBs, as dual loss leads to synthetic lethality^25,31^. It has previously been shown that this dependency can be rescued by *TP53* deficiency^31^. Our results confirmed this in a different cellular context, underscoring the importance of theta-mediated end joining in NHEJ/*p53*-deficient contexts. We also found that Pol θ inhibition reduced the recovery of bait-captured junctions mapping to RDCs in XRCC4/*p53*-deficient neural progenitor cells (Figure 4). Because LAM-HTGTS quantifies junctions to a defined bait (i.e., recoverable, joinable ends), this decrease is consistent with reduced translocation capture reported in Pol θ-or Ku80-deficient fibroblasts^26^. In addition, while our LAM-HTGTS data define a genotype-dependent role for Pol θ in RDC-derived end joining, they do not establish whether Pol θ is necessary or sufficient for CNV formation. The dependence of Pol θ in aphidicolin-induced CNVs was reported by Wilson et al. in the context of XRCC4-proficient human cell lines^8^.

We speculate that the reduced translocation efficiency we observe is linked to impaired microhomology-mediated end joining (Figure 4). This supports the role of Pol θ’s polymerase activity in promoting microhomology-dependent repair, which is consistent with previous literature^27,38^. We also observed a ∼15-base shortening of bait-derived DNA fragments (Figure 4), closely aligning with the gap-filling capacity of Pol θ observed *in vitro*^27^. These findings suggest that Pol θ may be required either to convert single-stranded DNA ends into double-strand breaks, or that ART558-mediated inhibition disrupts Pol θ-driven microhomology stabilization.

### The polymerase activity of Pol θ inhibits RDC end joining in wild-type neural stem and progenitor cells

In wild-type neural stem/progenitor cells, Pol θ inhibition led to increased RDC DSB end joining, in contrast to the trend observed in XRCC4/*p53*-deficient cells. We observed a marked increase in LAM-HTGTS captured DSBs, such as the *Lsamp*, *Grid2*, and *Ccser1* gene loci, implying that Pol θ operates to maintain genome stability at these RDC loci (Figure 5). Although striking, this finding is consistent with prior knowledge. We speculate that Pol θ protects DNA replication forks, thereby limiting the availability of RDC-derived ends for translocation capture. In this context, Pol θ has been shown to influence replication timing at a subset of genomic loci^28^, suggesting it may modulate the replication program in a context-dependent manner. Moreover, Pol θ protects genome stability during S phase and has been implicated in suppressing replication-associated damage outside G2/M phases^39^. In line with this, Pol θ knockout mouse embryonic fibroblasts exhibited nearly double the number of translocation events compared to *Ku80*/*Polq* double knockout cells^26^. Taken together, our data suggest that Pol θ constrains RDC-derived junction capture in wild-type neural progenitor cells by safeguarding genome stability during DNA replication.

### Limitation of the study

Given that RDCs are DNA replication-dependent, these findings suggest that CNV formation is initiated during S phase. However, due to limitations in our experimental setup, we were unable to directly determine whether CNVs are formed during the S phase or in the G2/M phase, as recently demonstrated ^8^. Nonetheless, we cannot exclude the possibility that a subset of S-phase-dependent RDCs persist into the G2/M phase, where they may be subsequently ligated. In addition, our methods could not resolve CNV frequency or measure CNV junctions in single cells. Further investigation using single-cell genomics to resolve CNV range and frequency would provide stronger support for linking our findings to clinical observations.

### Implications in neuronal disorders and cancer

Somatic CNVs in the brain have been widely linked to both neuropsychiatric disorders and cancer^5,40^. It has been hypothesised that CNVs may disrupt synapse formation by generating novel protein isoforms or deleting critical gene segments^18^. Given that many RDC-containing genes encode proteins involved in synaptic function, the resulting genetic heterogeneity, particularly in cell adhesion molecules, could impair the establishment of stable synaptic interactions.^41^. In addition, replication stress-induced CNVs have been previously proposed as a key driver of structural chromosomal instability in tumors^42^. For instance, the RDC containing genes: *LRP1B*, *NPAS3*, *LSAMP*, and *SMYD3* have all been found to be frequently deleted in glioblastoma^43,44^. Furthermore, large-scale CNV signature analyses across human cancers identified specific CNV patterns related to replication stress^5^. Notably, signature CN9, characterized by loss of heterozygosity and linked to TP53 mutations in glioblastoma, may arise through a similar replication stress-dependent mechanism reported by us. Our study provides a mechanistic basis that could help explain how such CNV signatures emerge under conditions of replication stress, with broader implications for tumor evolution.

## Material and Methods

### Mouse Embryonic Stem (ES) cell-derived neural progenitor cells (NPC)

Mouse embryonic stem (ES) cell-derived neural progenitor cells (NPCs) deficient in both *Xrcc4* and *p53* (*Xrcc4-/-p53-/-*) were used in this study, as previously described (Tena et al., 2020). The *Xrcc4-/-p53-/-* ES cells were cultured in DMEM supplemented with 15% ES cell-qualified fetal bovine serum, 20 mM HEPES, non-essential amino acids, 100 U/ml penicillin-streptomycin, 2 mM glutamine, 0.1 mM β-mercaptoethanol, and 1000 U/ml ESGRO recombinant mouse leukemia inhibitory factor (LIF, Millipore ESG1107). The cultures were maintained on a monolayer of irradiated mouse embryonic fibroblasts.

Differentiation of the *Xrcc4-/-p53-/-* ES cells into NPCs was performed following previously established protocols^20,45^. Briefly, the ES cells were seeded onto plates coated with laminin and poly-L-ornithine (Sigma Alderich, P4967) and cultured in N2B27 medium, consisting of 50% DMEM/F12 (ThermoFisher Scientific, 11330057), 50% Neurobasal (ThermoFisher Scientific, 21103049), 1% modified N2 supplement (Gibco, 17502048), 2% B27 supplement without retinoic acid (ThermoFisher Scientific, 12587-010), and 1× Glutamax (ThermoFisher Scientific, 35050061) for seven days. Subsequently, the cells were passaged onto laminin-coated plates and maintained for an additional five to six days in NBBG medium. This medium consisted of Neurobasal-A (ThermoFisher Scientific, 10888-022) supplemented with 2% B27 without retinoic acid, 0.5 mM Glutamax, 10 ng/ml human epidermal growth factor (EGF, ThermoFisher Scientific, PHG0314), and 10 ng/ml mouse basic fibroblast growth factor (FGFb, ThermoFisher Scientific, PMG0034).

### Primary Neural Stem and Progenitor Cell culture

The organ and embryo collection were covered under an institutional internal animal license (DKFZ381). Pregnant C57BL/6J female mice were sacrificed after 17 days of a positive vaginal plug. The embryonic (E) 17.5 embryos were retrieved, decapitated, and the brains were isolated for single cell suspension. In brief, brains were chopped into 2mm pieces, followed by papain digestion. Single cells were passed through a percoll gradient, washed with DPBS and NBBG, and were plated in the density of 1 million per mL in ultra low-attachment six-well plates (Merck, CLS3471). Fresh growth factors (EGF, FGFb, as described above, and PDGF, ThermoFisher Scientific, PMG0044) were added to the culture medium every second day. Five days after seeding, the neural spheroids were dissociated with TrypLE select (ThermoFisherScientific, 12563029), washed in DPBS and NBBG, and seeded to fresh ultra-low-attachment six-well plates in the same cell density. Five days after seeding, we dissociated the cells and performed nucleofection as described in the LAM-HTGTS section below.

### Whole genome sequencing

#### Library preparation

Approximately 5 × 10^6^ *Xrcc4-/-p53-/-* ES cell-derived NPCs were seeded onto freshly prepared poly-L-ornithine and laminin-coated plates and subsequently treated with either DMSO or 0.5 µM APH for a total of 96 hours. The treatment medium was refreshed after the initial 48 hours. Following treatment, cells were harvested, and genomic DNA (gDNA) was extracted with a standard phenol/chloroform/isopropyl protocol. Libraries were prepared following a TruSeq Nano DNA protocol by the NGS core facility staff at the DKFZ. Libraries were sequenced using an Illumina NovaSeq 6000 platform with 150 bp paired-end reads, achieving an average sequencing depth ranging between 160× and 200× per sample.

#### Alignment, QC, and CNV calling

Post sequencing, the raw FASTQ reads were subjected to automated AlignmentAndQC workflow (https://github.com/DKFZ-ODCF/AlignmentAndQCWorkflows/) aligned to the mm10 genome build. log₂(APH/DMSO) values were calculated using deepTools’ *bamCompare* function. To account for differences in sequencing depth between Aph– and DMSO-treated samples, we applied the built-in scale factor option. The accuracy of this normalization was verified by independently calculating the scale factor as the ratio of the average sequencing depth of APH to that of DMSO.

To quantify copy number variation across RDC-containing genomic regions, log₂ copy number ratio data log₂(Aph/DMSO) were analyzed using a custom R script. The script imported log₂ ratio data in bedGraph format and genomic regions of interest (in this case RDCs) in BED format. Non-standard chromosomes were filtered out, and the data were converted into GRanges objects. Overlapping bins between the log₂ ratio dataset and RDC regions were identified using the findOverlaps() function from the *GenomicRanges* package. For each RDC region, the average log₂ ratio was calculated across all overlapping bins. Results were exported as a tab-delimited text file.

To assess statistical significance, 10 sets of shuffled RDC regions were generated. These shuffled regions maintained the original lengths of the RDCs but were randomly repositioned across the mouse genome (mm10), ensuring chromosome sizes were respected. Each shuffled set was saved as a separate BED file for downstream analysis.

We also applied Delly2 for CNV calling. Four customized scripts were used for this purpose. Briefly, CNVs were identified from whole-genome sequencing data of Aph-treated and DMSO-treated samples. Read-depth profiles were generated for both Aph and matched DMSO BAM files, followed by SCNA calling using the mm10 reference genome and a predefined mappability blacklist. Somatic copy number alteration (SCNA) calls were merged and filtered for APH-specific events based on sample classification (Aph vs. DMSO), using thresholds including segment size ≥ 1,000 bp and ploidy level ≥ 2. Segmentation files were extracted and visualized. SVs were jointly called from Aph and DMSO BAMs, applying an exclusion list to mask problematic regions, and filtered for Aph-specific variants with a minimum mapping quality of 50. To improve somatic specificity, pre-identified variant sites were genotyped across a panel of unrelated DMSO controls, and Aph-specific SVs were retained based on consistent classification using updated sample metadata. The final filtered variants were annotated against the mouse mm10 gene model using a GTF annotation file.

#### Generating promoter-enhanced deleted cell lines for the *Ctnna2* and *Nrg3* loci

To generate *Ctnna2* and *Nrg3* promoter-enhancer-deleted (pe-del) ES cell clones, CRISPR/Cas9-mediated genome editing was performed using pairs of single guides RNAs (sgRNAs) designed to flank target regions. For *Ctnna2*, sgRNAs were positioned upstream of the gene’s *cis*-regulatory element and downstream of the last exon to create *Ctnna2*-deleted clones and flanking the entire *cis*-regulatory region to generate pe-del clones. For *Nrg3*, two sgRNAs were selected—one upstream and one downstream of the cis-regulatory elements—based on design and efficiency evaluation using ChopChop^46^. In both cases, cis-regulatory elements were defined based on histone modification marks (H3K4me, H3K9ac, H3K27ac) using ENCODE 3 ChIP-seq data from the UCSD/Ren lab. Individual ES cell clones were isolated, expanded, and screened by genotyping PCR to identify those carrying the intended deletions. Positive clones were further validated by Sanger sequencing to confirm precise deletion events.

#### Two-fraction replication sequencing (Repli-Seq)

Two fractions of Repli-seq were generated and analyzed as described^47^. In brief, ES cell-derived neural progenitor cells were plated in 30% confluence three days before BrdU incorporation. Cells were treated with 100uM BrdU (Sigma, B5002) for 2 hours, and cells were washed with cold DPBS ten times to remove loosely attached and dead cells. Cells were detached with Accutase, spun down, and resuspended in 2.5 ml ice-cold PBS plus 1% (vol/vol) FBS. Cells were fixed by adding 7.5 ml ice-cold 100% ethanol dropwise. The fixed cells were either stored at –20°C until sorting or directly stained with propidium iodide. Cell pellets were washed with ice-cold 1% (vol/vol) FBS in PBS once and resuspended in 0.5 ml of PBS/1% (vol/vol) FBS/propidium iodide (Sigma-Aldrich, P4170)/RNase A (Thermo Scientific, EN0531) solution as described. The final cell suspension concentration should be 4 × 10^6^ cells/ml. Cells were filtered through a 35 µm nylon mesh and directly to FACS. During the sorting, early or late replicating ES cell-derived NPCs were separated into two fractions; for each early and late fraction, 120,000 cells were sorted. The cells were pelleted, and genomic DNA was extracted as described. DNA was fragmented with a Covaris S220 Focused-ultrasonicator to 200-bp average fragment size. Libraries were constructed using the NEBNext Ultra DNA Library Prep Kit (New England BioLabs, E7370L) before BrdU pulldown, indexing, and amplification. Before submission, the libraries were further purified with AMPure XP beads (Beckman Coulter, A63881) and underwent quality control. After pooling, libraries were sequenced on Illumina Nextseq (75 bp single-read) or NovaSeq X-plus (100bp paired-read). After sequencing, the adaptors were trimmed from the raw FASTQ reads, and the reads were aligned to mm10/GRCm38 through Bowtie2 using a singularity container (https://github.com/brainbreaks/HighRes_RepliSeq). Underreplication score was determined by the Repli-Seq R package (https://github.com/CL-CHEN-Lab/RepliSeq) under the calculateURI function^14^, with the two-fraction Repli-Seq datasets.

### Linear amplification-mediated, High-throughput, genome-wide translocation sequencing (LAM-HTGTS)

#### Library used in this article

For comparing RDC and CNV positions (Figure 2), we extracted the published data (GSE233842) generated from the *Xrcc4-/-p53-/-* ES cell derived NPCs. We produced 89 primary LAM-HTGTS libraries for experiments shown for Figures 3-5, which were deposited at the GEO repository (GSE305347).

#### Library preparation

To induce targeted “bait” sites on chromosomes 6, 8, 12, or 14 in *Xrcc4-/-p53-/-* ES cell-derived neural progenitor cells or wild-type NSPCs, 5 × 10^6^ cells were nucleofected with 5 μg of SpCas9/sgRNA expression plasmids (pX330-U6-Chimeric_BB-CBh-hSpCas9; Addgene plasmid #42230). SgRNA sequences specific to each bait site were individually cloned into pX330 vectors following the protocol described by the Zhang lab (https://www.addgene.org/crispr/zhang/). Nucleofection was performed using the Nucleofector 2b device (Lonza) with the Mouse NSC Nucleofector Kit (VPG-1004, Lonza) according to the manufacturer’s instructions.

Following nucleofection, cells were cultured for 96 hours under treatments. For aphidicolin-treated conditions, cells were treated with 0.5 uM Genomic DNA was then extracted, and sequencing libraries were generated using established protocols. Libraries were sequenced on an Illumina NextSeq platform with the 150 pair-end chemistry.

#### Demultiplexing and alignment

Sequencing reads from demultiplexed FASTQ files were processed using the HTGTS Docker container (https://github.com/brainbreaks/HTGTS). The *TranslocProcess* module was used to trim adapters and perform library-specific demultiplexing. Trimmed reads were then aligned to the mm10/GRCm38 mouse reference genome using Bowtie2 and further processed using the *TranslocWrapper* function integrated into the pipeline. The file *result.tlx containing the bait and the prey information for each library were used for downstream analysis.

### DSB density, micro-homology usage, and bait-length analyses

To calculate the number of DSB to bait junctions detected in pre-specified regions-of-interest, we extracted the DSB coordinates from the *result.tlx files. All junctions were used to plot the junction density in Figure 2. For calculating the DSB density in the following figures, only DSBs identified at the non-bait chromosomes were considered. We used bedtools to count the number of recovered prey junctions per RDC in each LAM-HTGTS library. The number of recovered junctions per RDC was normalized using the total non-bait chromosomal DSBs, multiplied by a thousand, and divided by the length of RDC, in Megabase. This calculation derives DSB density as Junction per Thousand per Megabase, which is shown for Figures 4 and 5.

To calculate microhomology (MH) usage, we extracted information from the LAM-HTGTS outputfiles. We extracted *B_Qend*, which represents the length of the bait which translocates to an endogenous DSB, and *Qstart*, which represents the position where the endogenous DSB starts on the same read. Translocation junctions containing microhomology between 0-10 bp were kept for the calculation. MH was shown as frequency for Figs. 5 and 6. The length of MH was deduced by

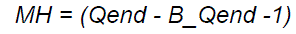

To calculate bait length (Blen), we calculate the distance between the beginning of the nested LAM-PCR primer (PrimStartCo) and the observed bait end or start (*B_Rend* or *B_Rstart* respectively) per read. All reads containing a junction were calculated, and the frequencies of bait lengths are shown for Figures 5-6.

In case of a bait DSB pointing at a *plus* orientation,

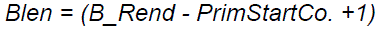

In case of a bait DSB pointing at a *minus* orientation,

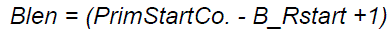

### Global run-on sequencing (GRO-seq)

GRO-seq libraries were prepared as described^16,24^. For each GRO-seq experiment, 5 – 10 million ES cell-derived neural progenitor cells’ nuclei were isolated for global run-on analyses. Two technical replicates per experiment condition were performed. GRO-seq FASTQ files were aligned to the genome build mm10/GRCm38 through Bowtie2 and processed using the GRO-seq pipeline (https://github.com/brainbreaks/GROseq).

### Reagents and laboratory equipment

Chemicals, cell culture reagents, solvents, primers and oligo sequences, antibodies, as well as specific laboratory equipment used for this article were presented in Supplementary Data 5.

## Acknowledgments

This work is supported by the Helmholtz Young Investigator grant, a Life and Health Alliance seed grant Program, and an ERC starting grant BrainBreaks to P-C W.. We thank Marco Giaisi for managing the Wei lab. We also thank the FACS core facility, the NGS core facility, the ODCF data management core facility, and the staff at the preclinic center at the DKFZ. We also thank the Körbel lab for fruitful discussion and suggestions. We extend our acknowledgements to Pavel Janscak and Jan Korbel for their valuable feedback. We also thank the team members at the Wei lab for intellectual discussions and technical support.

## Author Contributions

L.C. and P.-C. W. conceptualized and designed the project L. C., V. I., A. M., N. T., G. D. M., B. D., J. B., N.C. and M.G. performed the experiments: WGS (L.C., N.C.), ES cell pe-del clones (V. I., A. M., N. T.), Repli-Seq (L. C., B.D.), GRO-seq (L.C., V.I.), and LAM-HTGTS (L.C., G.D.M., B.D., M.G.). A. I., S. B., L. C., and P.-C. W. analyzed the data. L.C., A. I., V. I., A. M., and P.-C. W. interpreted the results. A. I., L. C., and P.-C. W. created figures and wrote the manuscript. T. H. supervised team member (S. B.). P.-C. W. secured research budget, supervised team members, and oversaw the research.

## Data Availability

The whole genome sequencing raw reads generated for this study were deposited in the European Genome Archive (PRJEB95922). Raw and processed files for two-fraction Repli-Seq, and LAM-HTGTS were deposited in the GEO (GSE305347). Raw and processed files for GRO-seq were deposited under GSE305346. Processed data were aligned to mm10 genome build. We retrieved high-resolution Repli-Seq data from GSE257765 (related to Figure 2). We also retrieved and LAM-HTGTS data from GSE233842 (related to Figure 2) and GSE74356 (wild-type Chr12 data, Figure 4; re-aligned to mm10) for replotting.

## Code availability

LAM-HTGTS, GRO-seq, Repli-seq pipelines are available at GitHub (https://github.com/brainbreaks/). Delly2 is a publicly available package (https://github.com/dellytools/delly).

## Supplementary figure legends

**Supplementary Figure 1.**
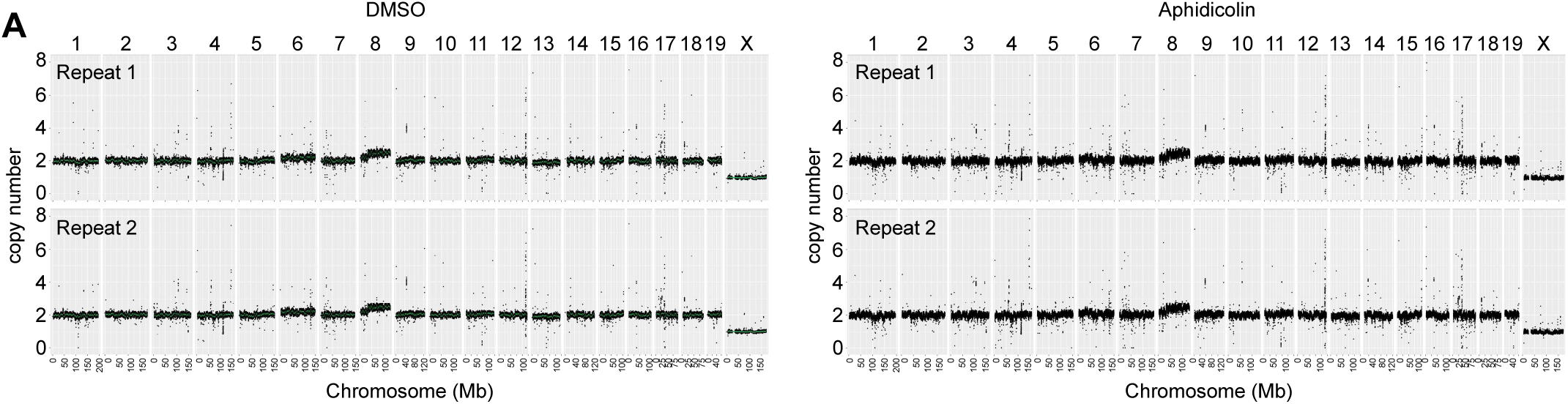
XRCC4/p53-deficient neural progenitor cell’s genomes were mostly diploid. The figure presents four ploidy plots for all autosomes and the X chromosome in samples treated with either the solvent control (DMSO) or aphidicolin. The Y-axis represents copy number, and the X-axis represents chromosome length in megabases.

**Supplementary Figure 2.**
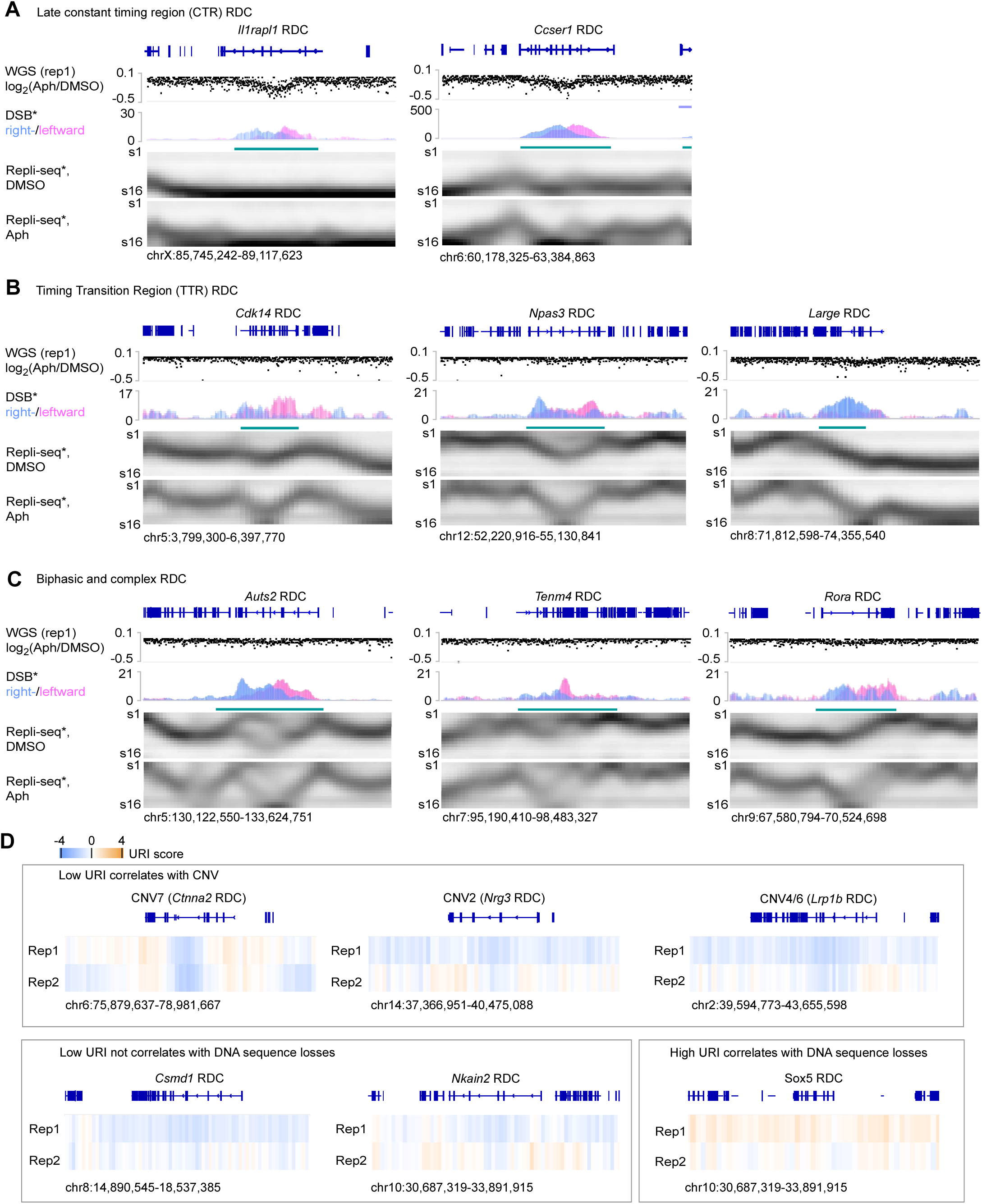
Aphidicolin-induced heterozygous deletions arise at a subset of late-replicating constant-timing RDC, continued. **(A)** Two RDC locations exhibited significant DNA sequence losses near the Delly2 cutoff. **(B)** Three RDCs are located at the timing transition region (TTR). **(C)** Three RDCs are located at the biphasic replicating region. Tracks in Panel A-C are as in Figure 2A. **(D)** Underreplication index (URI) and whole-genome sequencing log₂(APH/DMSO) for six RDC-containing genomic regions. Four loci—*Ctnna2*, *Lrp1b*, *Nrg3*, and *Csmd1*—exhibited low URI values and significant DNA sequence loss. At the *Sox5* locus, despite a high URI, a notable loss of DNA sequences was still observed. In contrast, the *Nkain2* locus displayed a low URI without corresponding DNA sequence loss. The heatmap scale is shown above.

**Supplementary Figure 3.**
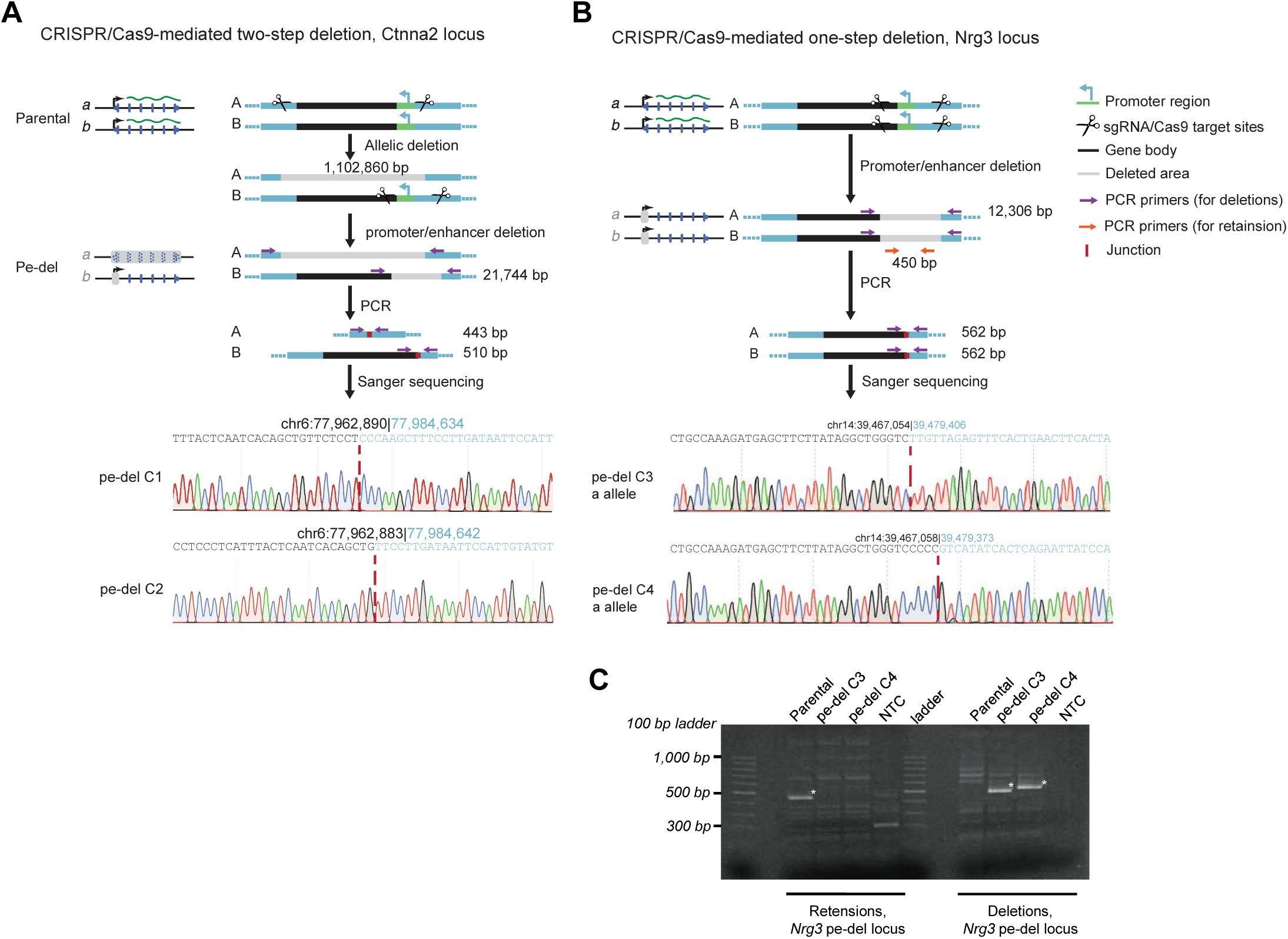
CRISPR/Cas9-mediated deletion strategies and validation for *Ctnna2* and *Nrg3* loci. **(A)** Two-step deletion strategy at the *Ctnna2* locus. Schematic of CRISPR/Cas9-mediated allelic deletion (step 1, 1,102,860 bp) followed by promoter/enhancer deletion (step 2, 21,744 bp). Parental configuration (top) and promoter/enhancer deletion (Pe-del) configuration (middle) are shown, with sgRNA/Cas9 target sites indicated by scissors and deleted regions in black. PCR primers for deletion (purple) and retention (orange) assays are indicated, with expected amplicon sizes noted. Sanger sequencing traces from independent *Ctnna2* Pe-del clones (C1 and C2) show the junction sequences, with target coordinates (mm10) labeled and the deletion breakpoints indicated by red dashed lines. **(B)** One-step deletion strategy at the *Nrg3* locus. Schematic of CRISPR/Cas9-mediated promoter/enhancer deletion (12,306 bp) in a single step. PCR primers and expected amplicon sizes are indicated. Sanger sequencing traces from *Nrg3* Pe-del clones (C3 and C4) show deletion junctions, with mm10 coordinates and breakpoints indicated as in (A). **(C)** PCR genotyping of *Nrg3* Pe-del clones. Agarose gel electrophoresis showing amplicons for retention (left panel) and deletion (right panel) assays in parental, Pe-del clones (C3 and C4), and none-template controls (NTC). Expected band sizes correspond to schematics in (B). Asterisks indicate the expected PCR products. As Sanger sequencing detected only one junction in either the C3 or C4 clones, these results indicate that both clones harbor a deletion on either the a or b allele extending beyond the regions targeted by the deletion-specific primers.

**Supplementary Figure 4.**
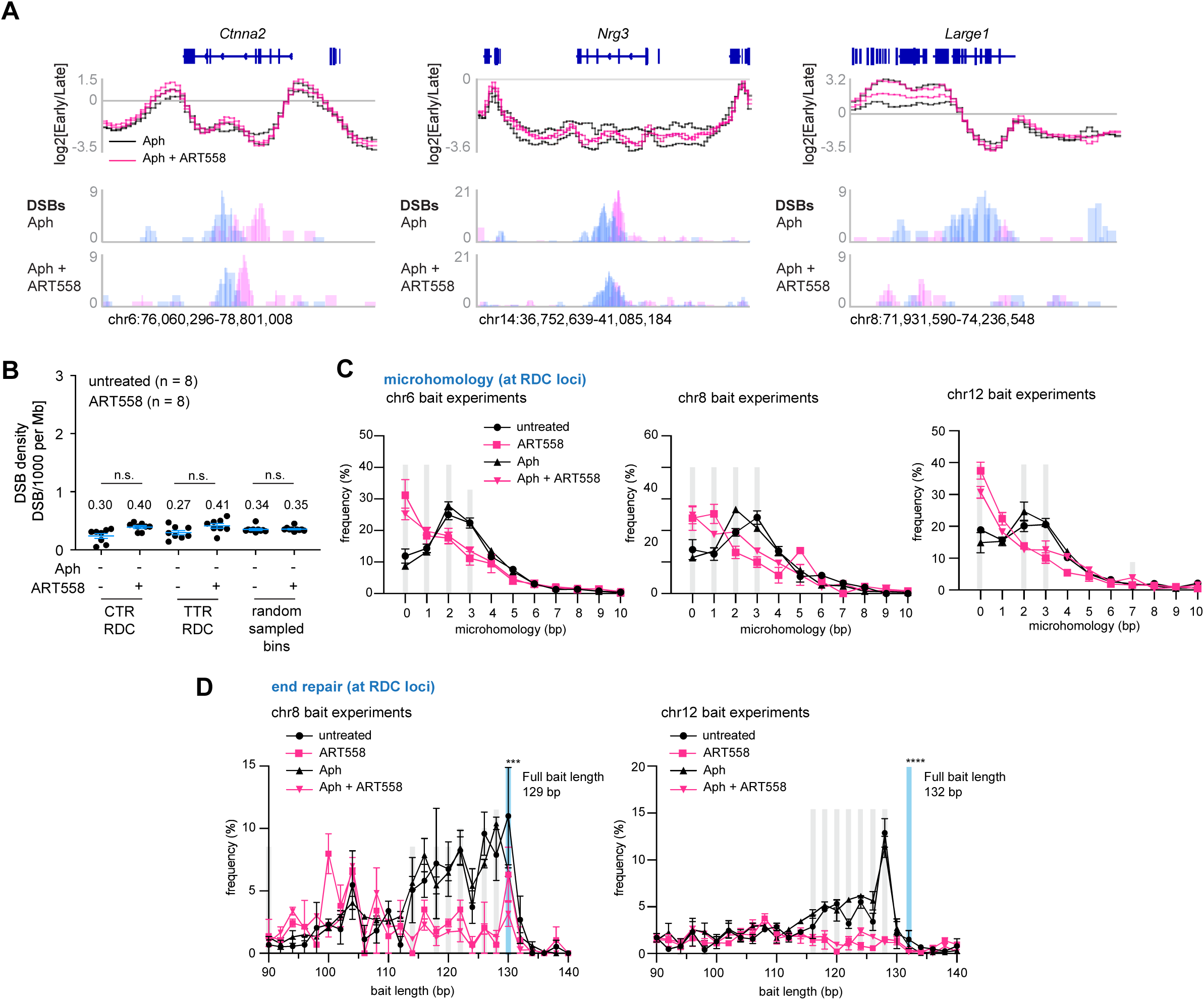
Pol θ inhibition re-wires the repair of replication-stress-induced DSBs. **(A)** Genome browser views of three large genes—*Ctnna2* (chromosome 6), *Nrg3* (chromosome 14), and *Large1* (chromosome 8). Upper traces: replication-timing profiles (Repli-seq, log₂ Early/Late) of cells exposed to low-dose aphidicolin (Aph, black) with or without the ATR kinase inhibitor ART558 (magenta). Lower histograms: LAM-HTGTS maps of DSBs obtained from the same samples (blue, Aph; pink, Aph + ART558). Grey horizontal lines mark the genome-wide mean. (**B)** Quantification of DSB density captured by LAM-HTGTS assays in 1520 randomly sampled equal-size genomic bins. Mean for each contusion was shown. Statistical power was determined by a two-tailed t-test. (**C)** Micro**-**homology frequency at RDC loci translocation junctions at chromosome 6, 8, and 12 baits. Mean and SEM were shown. (**D)** Distribution of recovered bait lengths in RDC-loci reads for chromosome 6 and chromosome 12 experiments. Mean and SEM were shown. The total length of bait without end resection is marked by a light blue column (164 bp for the chr14 bait). Bins showing statistically significant differences between APH and APH + ART558 samples, as determined by a two-tailed t-test, are indicated by grey shading. For **(C)** and **(D),** values for individual repeats are shown in Supplementary Tables 2-3; junction numbers analysed per experiment, and statistical significance per bin are shown in Supplementary Table 4.

**Supplementary Figure 5.**
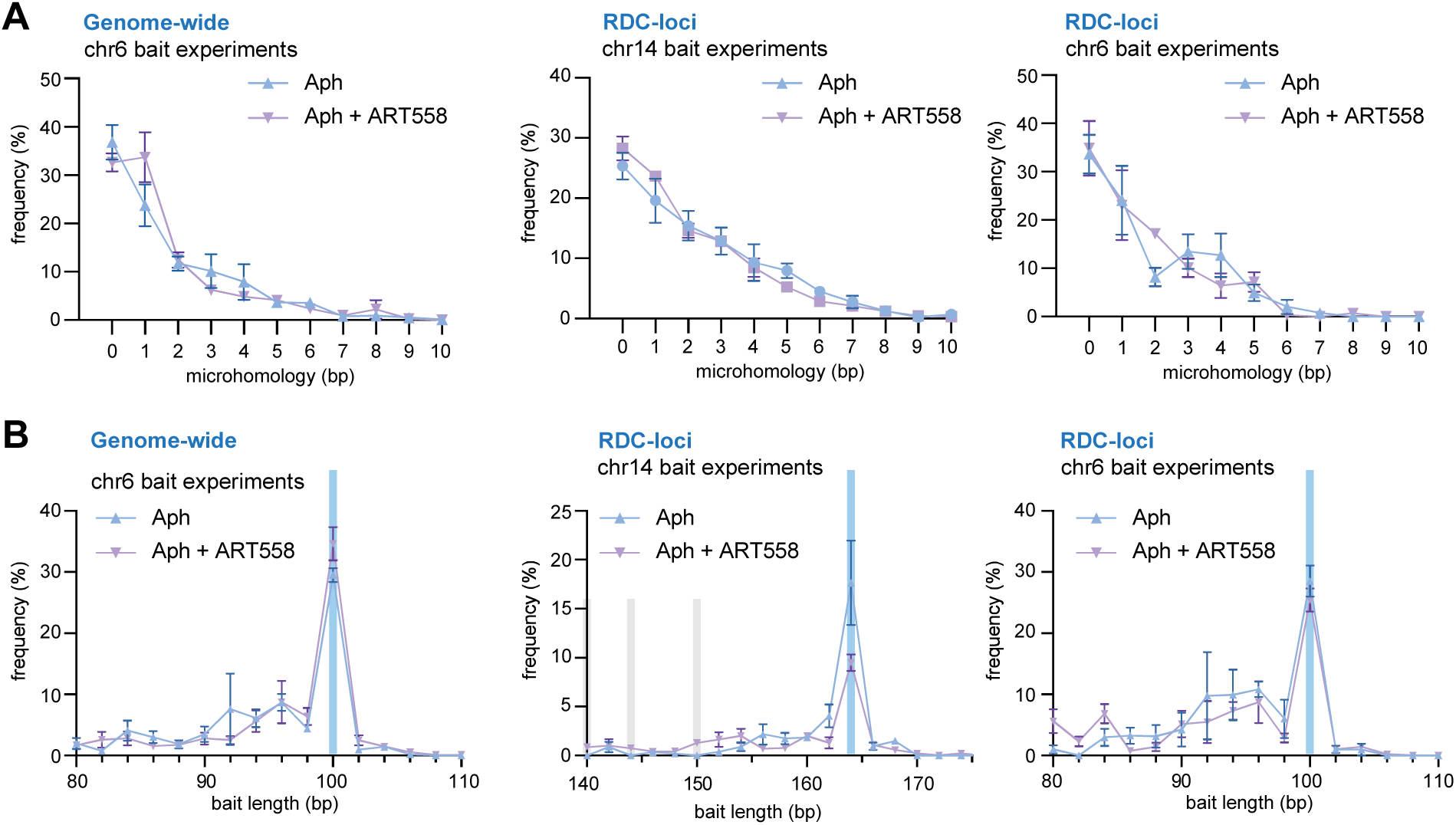
Pol θ inhibition slightly enhances direct joining, chromosome 6 experiments. **(A)** Micro**-**homology frequency at genome-wide translocation junctions at chromosome 6 baits, and junctions at RDC-loci for chromosome 6 and 14 baits. The figures show mean (dots) and SEM (error bars). Bins showing significant differences between two treatments are highlighted in grey. Statistical significance was determined using a two-tailed t-test. **(B)** Distribution of recovered bait lengths in deletion-sector reads for chromosome 6 and junctions at RDC-loci for chromosome 6 and 14 baits. An overview of bins 80 – 110 bp (chr6) and 140 – 170 bp (chr14) were shown. The figures show the mean (dots) and SEM (error bars). The total length of bait without end resection is marked by a light blue column (100 bp for the chr6 bait). Statistical significance was determined using a two-tailed t-test, and bins showing significant changes are highlighted in grey. Values for individual repeats are shown in Supplementary Tables 2-3; junction numbers analysed per experiment, and statistical significance per bin are shown in Supplementary Table 4.

**Supplementary Figure 6.**
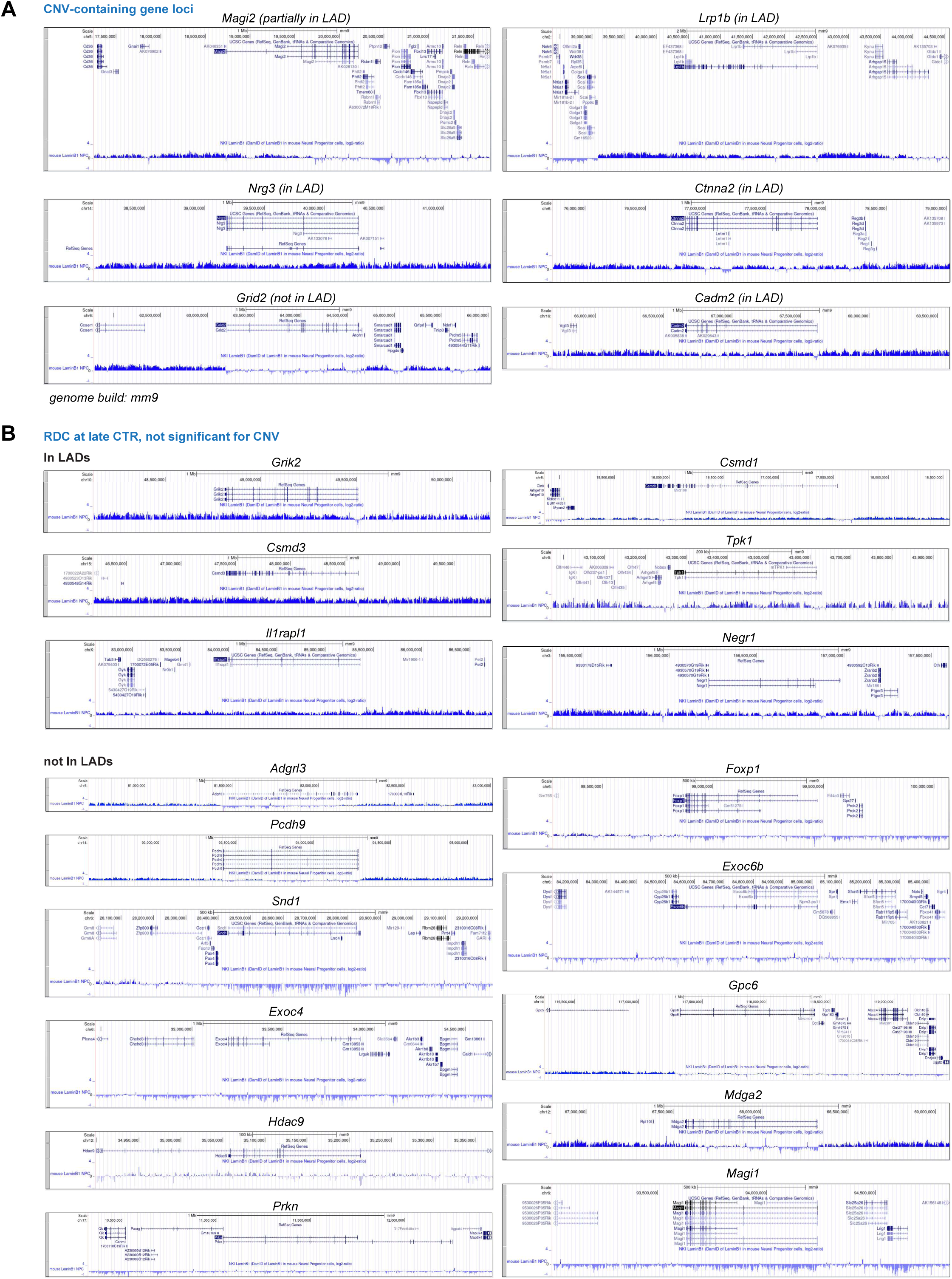
Most CNVs are embedded in the Lamine associated domain (LAD). (**A**) Genomic view of CNV loci located either within (*Magi2, Nrg3, Lrp1b, Ctnna2, Cadm2*) or outside (Grid2) LADs. The UCSC Genome Browser tracks show annotated genes and CNV-containing regions. Lamin B1 DamID signals in mouse neural progenitor cells are plotted as log2 ratios, with positive values indicating lamina association. **(B)** Genomic view of 16 representative late-replicating RDC-containing loci within (upper) or outside (lower) of LADs. Panels are shown as in (A). Lamin B1 DamID data were generated using an array-based approach and mapped to the mm9 genome assembly.

**Supplementary Data 1.** This table provides the coordinates and copy numbers for significant CNV loci shown for Figure 1 and 3. CHROM: chromosome, START: start of CNV, END: end of CNV, Copy Number APH: copy number in given CNV locus in cells treated with aphidicolin, and Gene: Gene overlap with CNV.

**Supplementary Data 2.** This table summarizes the frequency of microhomology (MH) usage. It presents joining junctions recovered either from genome-wide translocations or from translocations specifically targeting RDC loci. “Chr” denotes the chromosomal bait used in each experiment. MH (bp): microhomology in base-pairs, APH: aphidicolin, ART558: Pol θ inhibitor, and untreated: cells without APH of ART558 treatment.

**Supplementary Data 3.** This table summarizes the frequency of bait length (bp) for reads involving end-joining. It presents joining junctions recovered either from genome-wide translocations or from translocations specifically targeting RDC loci. “Chr” denotes the chromosomal bait used in each experiment. Blen (bp): bait length in base-pairs, APH: aphidicolin, ART558: Pol θ inhibitor, and untreated: cells without APH of ART558 treatment.

**Supplementary Data 4.** This table provides numbers of junctions analyzed for Figures 4 and 5 experiments. It also contains statistical significance values for results shown for Figures 4 and 5.

**Supplementary Data 5.** This table provides information for reagents, oligonucleotides, and equipment used for this article. Vender, catalog number, and purposes of assay for each reagent were shown. All reagents are commercially available.

